# A Family of Interaction-Adjusted Indices of Community Similarity

**DOI:** 10.1101/040097

**Authors:** Thomas Sebastian Benedikt Schmidt, João Frederico Matias Rodrigues, Christian von Mering

## Abstract

Interactions between taxa are essential drivers of ecological community structure and dynamics, but they are not taken into account by traditional indices of diversity. In this study, we propose a novel family of indices that quantify community similarity in the context of taxa interaction networks. Using publicly available datasets, we assess the performance of two specific indices which are *Taxa INteraction-Adjusted* (TINA, based on taxa co-occurrence networks), and *Phylogenetic INteraction-Adjusted* (PINA, based on phylogenetic similarities). TINA and PINA outperformed traditional indices when partitioning human-associated microbial communities according to habitat, even for extremely downsampled datasets, and when organising ocean micro-eukaryotic plankton diversity according to geographical and physicochemical gradients. We argue that interaction-adjusted indices capture novel aspects of diversity outside the scope of traditional approaches, highlighting the biological significance of ecological association networks in the interpretation of community similarity.

## Introduction

Understanding how patterns of diversity are established and maintained is fundamental to the ecological characterisation of living systems. Following Whittaker ^1,2^, *diversity* is traditionally considered to comprise three components: local diversity of individual habitats *(α diversity)* and between-site variation (β *diversity*) together determine the total diversity of a given system *(γ diversity)*. However, while referring to these definitions, researchers have studied conceptually different phenomena under the umbrella term “diversity”. β diversity, in particular, has been variously reported as species turnover or variation, further sub-defined and quantified using different mathematical approaches ^3,4^. Nevertheless, most authors agree that *community similarity*, the compositional variation between sites, is an integral aspect of β diversity, and more generally one of the most important parameters in community ecology ^5^. To characterise the mechanisms underlying an observed diversity structure, it is essential to quantify and appraise patterns of community similarity.

A multitude of mathematical indices of community similarity have been proposed: as of 2015, the widely used software *EstimateS* ^6^ computes 16 different indices, while the popular microbial ecology toolboxes mothur ^7^ and phyloseq ^8^ provide as many as 37 and 44 measures, respectively. The various available measures capture conceptually different aspects of diversity. Traditional indices, such as the Jaccard ^9^ or Bray-Curtis ^10^ index, focus on taxa compositional overlap, quantified directly from taxa count data. More recently, phylogenetically informed indices have become increasingly popular which, in contrast to census-based metrics, do not treat taxa independently but rather quantify shared evolutionary history between communities ^11,12^. Traditional and phylogenetic metrics may provide complementary insights on the processes driving community composition, particularly since phylogenetic relatedness of taxa is a proxy for functional or ecological similarity ^13^.

Apart from analysing diversity patterns, another important approach to characterising ecosystem function focuses on studying interaction networks of ecological or functional associations between taxa directly. Applying graph theory to food webs, mutualist or host-parasite networks and others has revealed an important role for interaction structure in community stability and dynamics ^14–16^. This approach has been particularly fruitful in microbial ecology, where “true” ecological interactions can to a certain extent be inferred from co-occurrence networks of anonymous *Operational Taxonomic Units* (OTUs)^17,18^. Highly informative taxa co-occurrence networks have been constructed for many ecological systems, including the human body-associated microbiota ^19,20^ or ocean planktonic communities ^21^, as well as for global, integrated datasets across various habitats ^22^.

One main difference between such diversity-based and network-based approaches lies in the analysis scope: the latter identify drivers of community structure at the level of individual taxa interactions, while the former reveal compositional patterns at community level. Arguably, both approaches are informative, but they are often pursued independently: it remains challenging to interpret community-level diversity changes in the light of taxa-level ecological associations, and vice versa. In this study, we propose to bridge this analysis gap by developing a set of mathematical indices that quantify community similarity (or β diversity) as the average taxa interaction strength between samples. While our method is applicable to many types of interaction data, we focus on *Taxa INteraction-Adjusted* indices (TINA), based on taxa co-occurrence data, and *Phylogenetic INteraction-Adjusted* indices (PINA), based on phylogenies. In a re-analysis of two publicly available datasets, we show that TINA and PINA capture known diversity patterns better than existing indices, even for very small datasets, and that they can reveal novel and refined biological interpretations.

## Methods

In this study, we compared a total of 11 indices of community similarity (listed in Table 1) that fall into three categories, “traditional” taxa count-based indices, phylogenetic indices and our proposed interaction-adjusted indices (Figure 1).

**Table 1.**
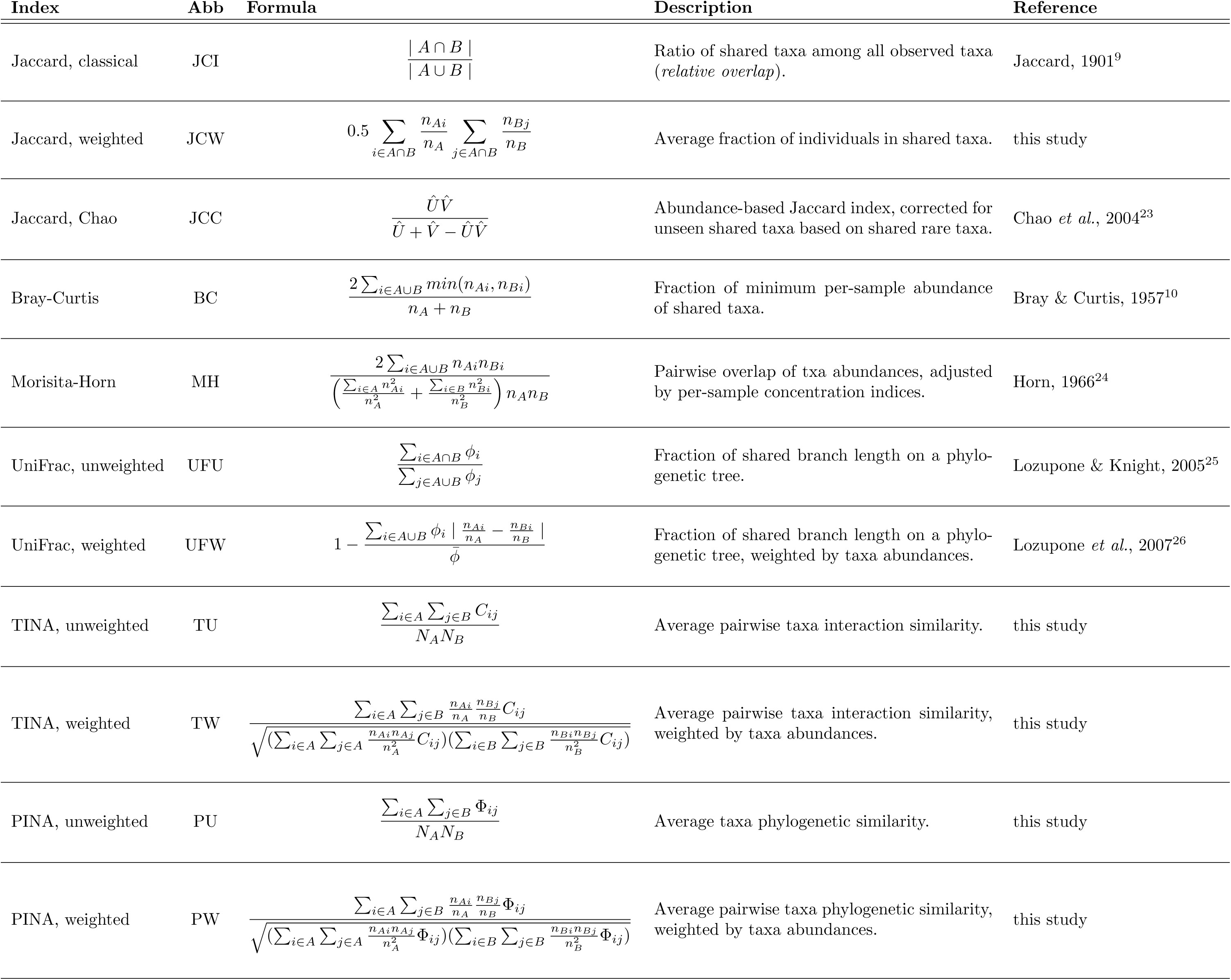
Overview of different indices of community similarity used in this study. All formulas and descriptions are given as similarities; for ordinations and statistical tests, corresponding distances or dissimilarities are used (D = 1 - S). N_A_, total number of taxa in sample A; n_A_ total number of individuals in sample A; n_Ai_, individuals of taxon i in sample A; Û, estimator for fraction of individuals in shared taxa for sample A, according to formula 9 in Chao et al. ^23^; φ_i_, phylogenetic branch length of taxon i to root; Φ_ij_, phylogenetic association between taxa i and j; C_ij_, interaction similarity between taxa i and j.

**Figure 1.**
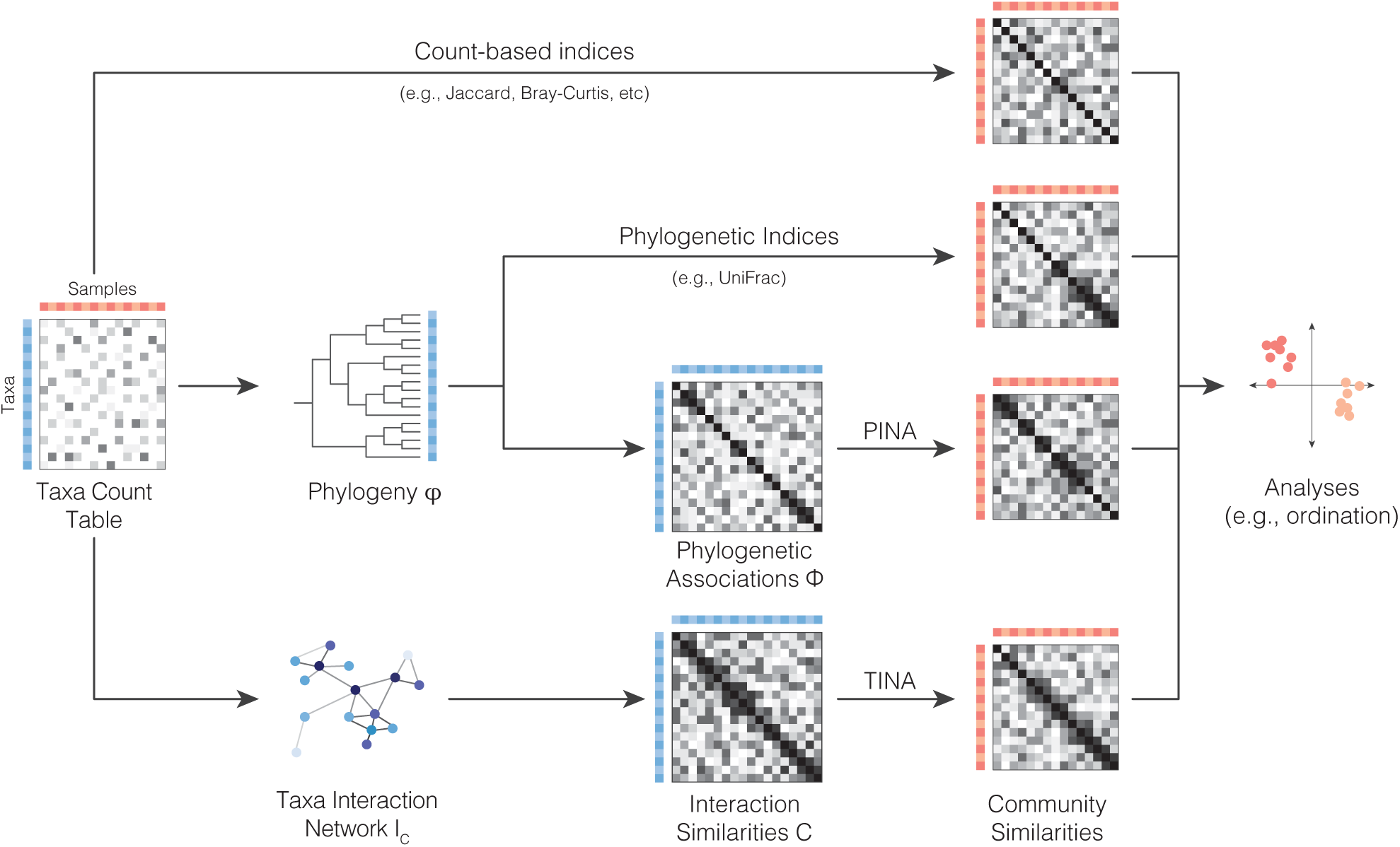
Overview of different approaches to quantifying community similarity. Based on a taxa-sample count table, traditional count-based indices such as Jaccard and Bray-Curtis quantify community similarity from the overlap in taxa composition (upper branch). In contrast, phylogenetic indices such as UniFrac take into account taxa relationships, quantifying community similarity as shared evolutionary history, based on taxa phylogeny (middle branch). Our proposed Taxa INteraction-Adjusted (TINA) and Phylogenetic INteraction-Adjusted (PINA) indices, in contrast, take into account similarities on a taxa co-occurrence network, codified in an interaction similarity matrix C, or in terms of cophenetic phylogenetic distances, represented in a phylogenetic association matrix Φ.

### Classical and phylogenetic community similarity indices

In his widely cited "*comparative study of the floral distribution of parts of the Alps and the Jura”*, Paul Jaccard ^9^ introduced what is arguably the earliest index of β diversity. For two communites A and B, the *classical Jaccard index* (JCI) is the relative taxa overlap, i.e. the ratio of shared taxa among all sampled taxa (see formula in Table 1). In this original fomulation, the JCI is *incidence-based*, or *unweighted*: it considers only the presence and absence of taxa, but not their relative abundance ratios. Several *abundance-based* or *weighted* variations of the classical Jaccard index have been proposed ^23^; here, we use a straightforward *weighted Jaccard* (JCW) formulation which describes community similarity as the mean fraction of individuals in shared taxa across both focal samples. While the JCI has proved very versatile and is used for manifold scientific problems beyond biology, an ecology-specific variant that corrects for the characteristics of imperfect sampling has been proposed by Chao et al. ^23^: *Chao’s weighted Jaccard* index (JCC) extrapolates the fractions of individuals in unseen shared taxa based on the number of observed rare taxa.

One of the most widely used indices in modern community ecology is arguably the *Bray-Curtis (dis)similarity* (BC) ^10^ which describes community overlap as the fractional minimum abundance of shared taxa between samples. Somewhat related to BC is the *Morisita-Horn overlap index* (MH) ^24^, calculated as pairwise multiplicative taxa overlap, adjusted by a per-sample concentration index.

These classical indices and their derivations assess community overlap directly from count data, treating all observed taxa as independent (Figure 1, top branch). Phylogenetic indices, in contrast, consider the (phylogenetic) relationships between taxa and quantify community similarity as shared evolutionary history (Figure 1, middle). Among these increasingly popular indices is *UniFrac* which calculates the shared branch length between samples on a phylogenetic tree, either for observed taxa based on incidence (unweighted UniFrac, UFU)^25^, or based on taxa abundances (weighted UniFrac, UFW)^26^.

### A novel family of Interaction-Adjusted community similarity indices

Consider two communities A and B, composed of N_A_ and N_B_ taxa from which a total of n_A_ and n_B_ individuals have been sampled. Next, consider a matrix I which describes pairwise taxa interactions, such that I_ij_ is the interaction between taxa i and j. Manifold types of interactions with different biological meanings, different layers of information and at different levels of curation effort are suitable, such as e.g. predator-prey relationships, symbiosis, parasitism, mutualism, cross-feeding, resource competition, etc. Here, we consider the case of ecological associations as inferred by taxa co-occurrence networks, constructed from taxa count tables by pairwise association of abundances across samples (Figure 1, bottom). The scale and characteristics of such a co-occurence interaction matrix I_C_ will depend on the association metric chosen; Faust & Raes ^18^ have provided a comprehensive review of different approaches to network construction and interpretation. For example, a taxa abundance correlation network would scale from −1 (avoidance) to +1 (complete association), while other popular association metrics may scale differently. Therefore, it is important to transform the interaction matrix I_C_ to a common scale; we do this by correlating taxa by their pairwise associations to all other taxa in the system (i.e., row I_C,i*_ of I_C_ for taxon i) and transforming this into a Pearson similarity:

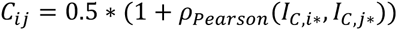

Thus defined, a transformed co-occurrence matrix C has several desirable properties: (i) it scales from 0 (avoidance) to 0.5 (neutral association) to 1 (complete association); (ii) C_ij_ corresponds to the “proximity” of taxa i and j on the original association network I_C_; (iii) the transformation generally sharpens and smoothens network structure, amplifying strong associations and down-weighting spurious correlations, but association ranks remain mostly unchanged (see also Figure S1 in Supporting Information).

Given this transformed interaction matrix, we propose to quantify the similarity between communities A and B as the *average interaction strength between all taxa observed in A or B*. We thus define an incidence-based or unweighted *Taxa INteraction-Adjusted* index of community similarity (*unweighted TINA*, TU) as

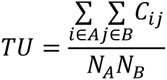

Likewise, an abundance-based or *weighted TINA* index (TW) can be defined as *weighted average taxa interaction strength*, scaled by the geometric mean per-sample weighted taxa interaction strength:

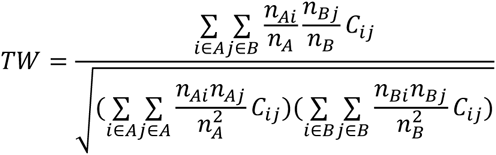

TINA values are 1 for two completely identical communities, but also if all taxa in A and B are perfectly associated. If no taxa are shared, TINA values tend towards 0.5 if taxa interactions are neutral (neither associative nor dissociative) and towards 0 if taxa between A and B show a strong avoidance signal. Thus, TINA resolves non-zero similarities even for pairs of communities that do not share any taxa (which implies zero similarity according to traditional, count-based indices); theoretically, the TINA index for such disparate pairs can even be 1 if all their taxa are perfectly associated.

TINA-like indices can be defined analogously for any kind of interaction data, given that interaction matrices are transformed similarly to the I_C_ to C transformation described above. This is also true for the special case of phylogenetic “interactions”. Consider a phylogenetic tree φ of taxa observed in a given system with a *cophenetic phylogenetic similarity* matrix I_φ_ which can be interpreted as a phylogenetic association network (analogous to I_C_) and transformed into an association matrix Φ (analogous to the co-occurrence association matrix C). Then, we can define *unweighted Phylogenetic INteraction-Adjusted* community similarity (*unweighted PINA*, PU) as

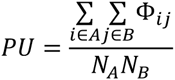
and *weighted PINA* (PW) as

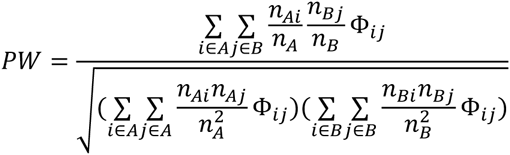

### Human Microbiome Project data analysis

To test the performance of different community similarity indices, we re-analysed two publicly available datasets, provided by the Human Microbiome Project and the TARA Oceans project, respectively. Raw 16S rRNA V35 region sequencing reads of the Human Microbiome Project ^27^ were downloaded from the NCBI Sequence Read Archive; metadata was obtained from the HMP data repository (hmpdacc.org). Sequences were filtered for chimeric reads using UCHIME ^28^, aligned to a secondary structure-aware 16S rRNA model using Infernal ^29^, denoised by a global minimum read abundance at 1% tolerance of 4 and clustered into OTUs at 97% average linkage sequence similarity using hpc-clust ^30^, as established previously ^31,32^. The resulting filtered taxa count table contained 24,717,447 sequences clustered into 27,041 OTUs across 3,849 samples. A phylogenetic tree of OTU representatives, selected by minimum average within-OTU sequence distance, was inferred using FastTree2 ^33^ with default parameters. Pairwise taxa correlation networks for the full dataset and subsets were calculated using a custom R implementation of SparCC ^34^, an adapted correlation metric correcting for spurious associations that has been shown to approximate “true” ecological interactions ^17^.

### TARA Oceans data processing and analysis

From the TARA Oceans eukaryotic plankton diversity census ^35^, we downloaded 18S rRNA V9 tag data organised into an OTU-level taxa count table (doi.pangaea.de/10.1594/PANGAEA.843022) and sample metadata (doi.pangaea.de/10.1594/PANGAEA.843017). Data per sample were pooled across filter sizes and OTUs containing ≤30 sequences and several orphan samples were removed (see analysis code), yielding a filtered count table of 535,903,407 sequences, 27,448 OTUs and 77 samples for which a SparCC correlation network was computed.

### Data and software availability

All analysis code and processed datasets are available online

(github.com/defleury/Schmidt_et_al_2016_community_similarity;

meringlab.org/suppdata/2016-community_similarity/).

## Results

### TINA and PINA provide improved partitioning of human body site-specific microbial communities

From an ecological point of view, the human body appears as little more than a system of distinct microbial habitats ^36^. The *Human Microbiome Project* ^27^ has provided a first comprehensive census of human body-associated microbial communities and their potential functional repertoires. HMP 16S rRNA tag sequencing data is available for 18 habitats from five different body sites, namely oral cavity (9 habitats), skin (4), female urogenital tract (3), airways (1) and gastrointestinal tract (1); see Figure 2 for a full list. One original goal of the HMP, similar to many ecological studies, was to establish how compositional similarity patterns distinguish communities associated to these different habitats. In other words: are body sites distinct from each other in microbial community composition, and which other factors drive compositional variance?

**Figure 2.**
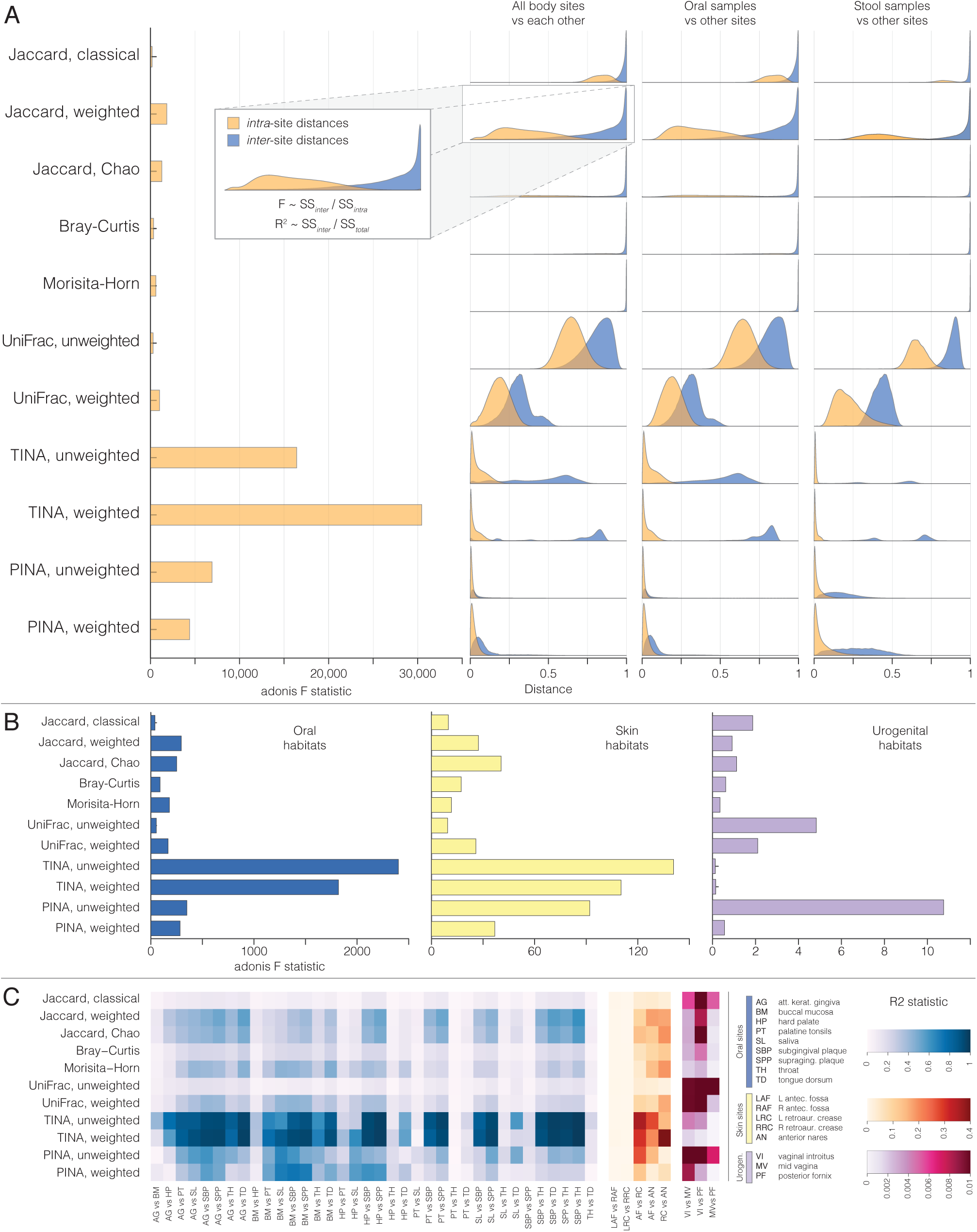
Differential partitioning of human body habitat-specific community structure. (A) Partitioning by general body sites. Left, PERMANOVA F statistics for different indices when testing community distance partitioning according to general body site, i.e. into oral, skin, urogenital, airways and gastrointestinal habitats. Right, histograms of community distances intra-site (orange) and inter-site (blue) for all body sites against each other, oral against other habitats and gastrointestinal against other habitats. The inset illustrates how PERMANOVA F statistics and R2 values are calculated from community distances using the *adonis* function of the R package *vegan* ^37,38^. (B) Sub-partitioning of oral habitats (blue), skin habitats (yellow) and urogenital habitats (violet). (C) Pairwise separation by different indices of all pairs of oral habitats, skin habitats against each other and against airways (anterior nares) and urogenital habitats against each other, as detected by the PERMANOVA R^2^ value (relative variance explained by factor habitat). Note the different color scales, indicating overall differential partitioning power per body site.

These types of questions are typically addressed by calculating pairwise community distances and then applying multivariate statistical tests to establish how much of the distance matrix’ structure is explained by a given model. One of the most widely used methods is Anderson’s PERMANOVA (permutational analysis of variance)^37^, implemented in the *adonis* function of the R package *vegan* ^38^, which calculates a pseudo-F statistic on group separation from the sums of squares of *inter*-group distances over the sums of squares of *intra*-group distances (see Figure 2A) and then conducts a permutational significance test. Thus, the adonis F statistic, as well as the related R^2^ statistic (the ratio of variance explained by the tested factor) indicate an *effect size* of multivariate group separation (higher F and R^2^ values indicate more discriminatory power), while a permutational p value indicates significance. F and R^2^ statistics have previously been used to benchmark multivariate ecological analyses, e.g. by Eren et al. ^39^.

We reanalysed the HMP data using 11 different community similarity indices (Table 1), five of which are count-based (JCI, JCW, JCC,BC and MH), two phylogenetic (UFU, UFW) and four interaction-adjusted (TU, TW, PU and PW). We observed that partitioning of the five general body sites (oral cavity, skin, urogenital tract, airways, gut) by community similarity was by far best for TINA (F=30,455 for TW; F=16,421 for TU) and PINA (PW, 4,397; PU, 6,917) when compared to all other indices, with JCI providing the weakest discrimination (F=182). This improved partitioning was due to several effects, as indicated by community distance histograms per index (Figure 2A, left). First, TINA and PINA provided very high overall resolution, distributing pairwise dissimilarities across a broader range on the interval 0 (identical communities) to 1 (complete dissimilarity) than most count-based indices. Second, TINA and PINA assigned very low and sharply distributed intra-group dissimilarities, meaning that samples from the same body sites were on average considered very similar to each other; in contrast, count-based indices showed very sharp and pronounced inter-group dissimilarities, but wider distributions within groups. Finally, intra- and inter-group dissimilarities were generally much clearer separated for TINA than for count-based indices or UniFrac.

### Combined interpretation of TINA and PINA may provide novel biological insights

While habitat partitioning was differentially pronounced, all indices provided significant group separation (p≤0.001, 999 permutations). Indeed, differences in community composition between body sites – which are highly distinct micro-environments – can be expected to be large, so it is not surprising that they were picked up by all indices. We therefore conducted similar tests on more complicated problems, such as the separation of habitats within a body site (Figure 2B) or of pairs of similar habitats (Figure 2C). TINA provided by far the strongest partitioning of oral and skin habitats, followed by PINA and JCW/JCC. For urogenital sites, in contrast, only few indices provided significant separation at all: unweighted PINA (PU), UFU, UFW and JCI. These trends were consistent with pairwise separability of habitats (Figure 2C) which was highest for TINA in oral and skin, but for PU, UFU and UFW in urogenital sites.

These observations imply that diversity patterns in these habitats are determined by different factors. TINA quantifies community similarity as an overlap in ecological associations of taxa, while PINA and UniFrac focus on shared phylogeny. Thus, it appears that the compositional identity of oral and skin sites is driven by recurring cliques of associated OTUs, as captured by strong co-occurrence signals, while pairwise taxa associations are less important in the urogenital tract, where communities of changing partners are instead filtered by phylogeny, possibly indicating a functional signal.

### TINA captures habitat structure of the human microbiome taxa co-occurrence network

To illustrate how TINA captures ecological taxa interaction structure, how TINA values can be interpreted at the level of individual sample pairs and under which conditions it provides more intuitive results than count-based indices, we selected 16 HMP samples for which Figure 3 shows the taxa co-occurrence network; Figure S2 shows the same network, coloured by OTU phylum-level taxonomy. We observed that the network is structured into several habitat-specific clusters of strongly co-occurring OTUs, with slightly less dependence on taxonomy. This is in line with the general observation that (microbial) co-occurrence networks tend to capture ecological signals which indeed can often be much more subtle than the present body habitat classification ^18^.

**Figure 3.**
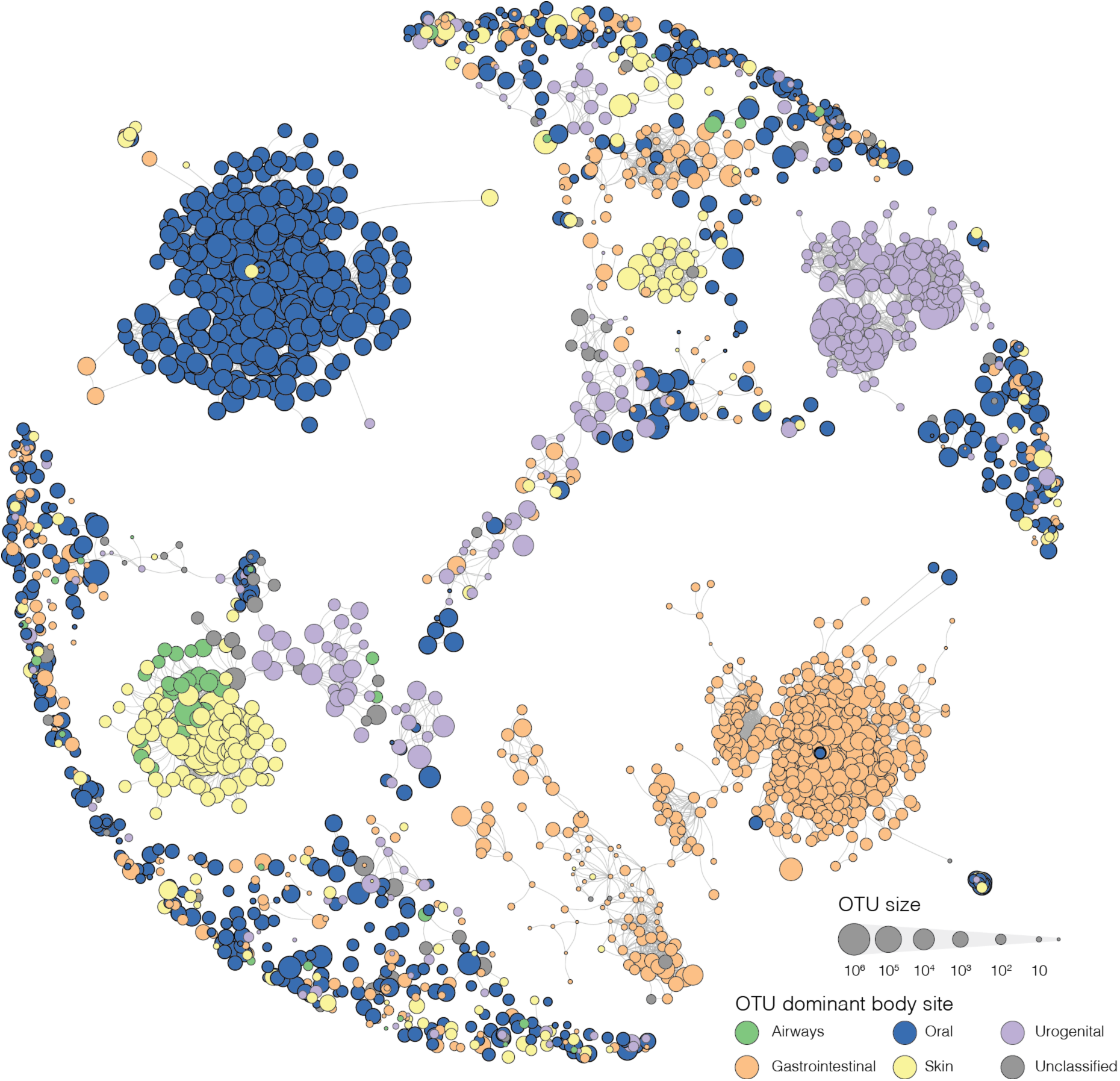
Taxa co-occurrence network for a subset of Human Microbiome Project samples. 16 samples from the full HMP dataset were selected as described in the main text, comprising a total of 2,671 OTUs for which all pairwise SparCC correlations ≥0.5 are shown as edges in the network. Node size indicates global OTU size, i.e. the total number of counts per OTU across the full HMP dataset. Node colour indicates OTU dominant habitat, assigned if more than 50% of all OTU abundance was in samples of the same body site. Figure S2 shows the same network, coloured by OTU phylum-level taxonomy.

Next, we mapped three examples of sample pairs onto this network to illustrate how TINA takes interaction structure into account to quantify diversity. In the first example (Figure 4A), we consider two samples from the vaginal posterior fornix which where the most similar pair according to the classical Jaccard index (lowest JCI distance). These samples have a high taxa overlap, while many of their non-shared taxa also share strong co-occurrence signals, so that TINA likewise assigns them a very high similarity. Thus, in this case, TINA and count-based indices agree in considering both samples highly similar.

**Figure 4.**
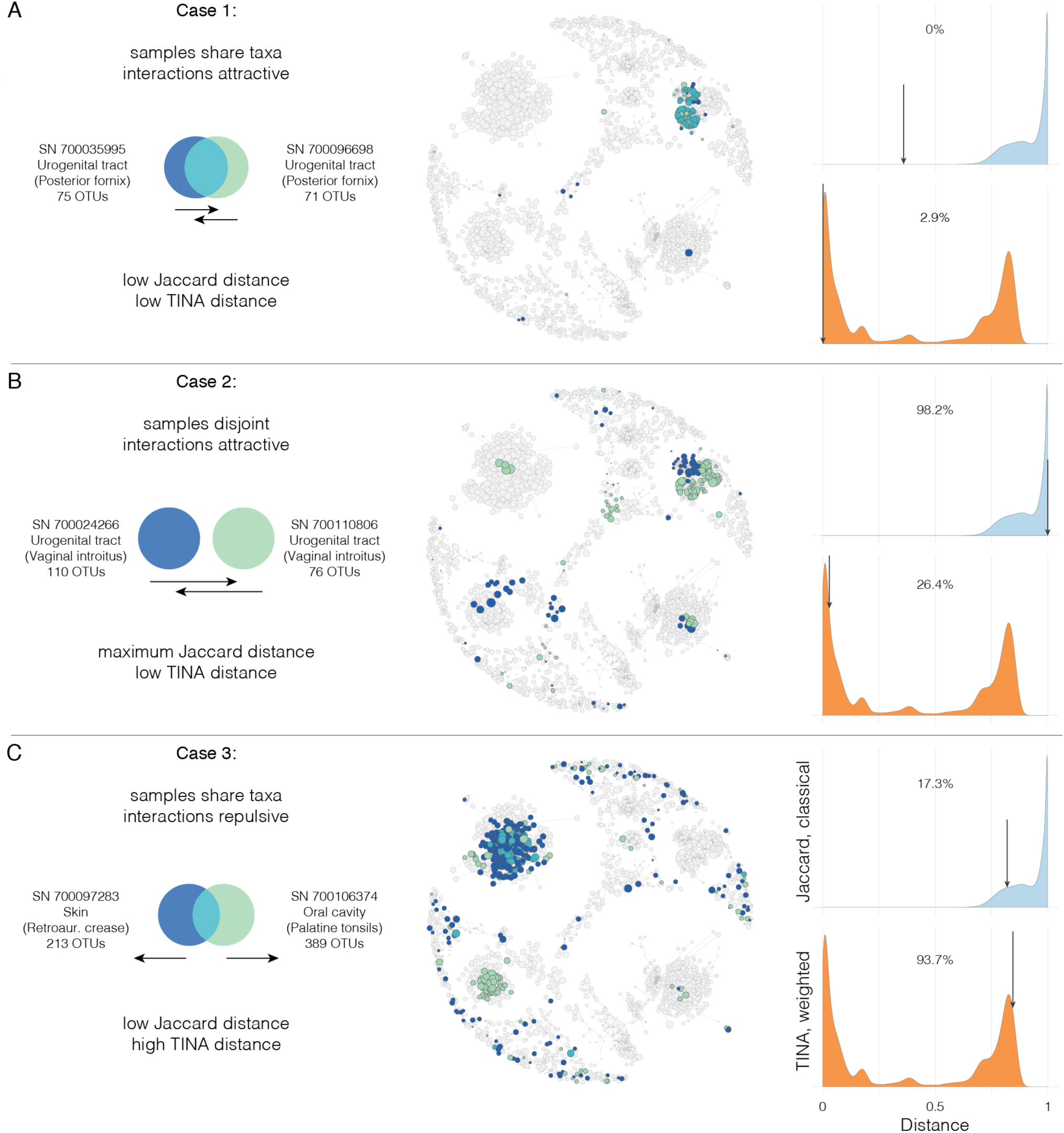
TINA quantifies community similarity from taxa co-occurrence. (A) For two urogenital samples which share a large taxa overlap, both traditional count-based indices (exemplified by the classical Jaccard index, JCI) and TINA assign a low-ranking community distance (as indicated in community distance histograms on the right). The middle panel shows how taxa of these samples map onto the co-occurrence network introduced in Figure 3; blue, taxa unique to sample SN700035995; green, taxa unique to sample SN700096698; blue-green, taxa shared between both samples. (B) For two urogenital samples that do not share any taxa, but whose taxa still share attractive co-occurrence interactions, JCI assigns complete distance (JCI=1), while TINA assigns a relatively low-ranking distance. (C) In the opposite case of two samples from different body sites (skin and oral) which have a significant taxa overlap, but repulsive taxa interactions, JCI assigns a low-ranking, but TINA a very high-ranking distance.

The second case is less trivial (Figure 4B). Here, we consider two urogenital samples from the vaginal introitus which do not share any taxa at all – count-based indices assign a distance of 1, considering them completely dissimilar. However, the taxa found in these samples share an attractive interaction signal, although some taxa form part of distinct co-occurence clusters. In this case, TINA provides a more intuitive result by ranking the similarity between two samples from the same habitat (vaginal introitus) relatively high.

Finally, consider the opposite situation (Figure 4C). Here, two samples from different body sites (skin and oral) happen to share several taxa, so that JCI ranks them among the top 17.3% most similar samples. However, most non-shared taxa belong to distinct co-occurrence clusters, meaning that their pairwise interactions are repulsive, such that TINA assigns a very high dissimilarity to this pair which, again, is an arguably more realistic picture.

### Interaction-Adjusted Indices provide strong partitioning even for small datasets

The HMP dataset used in this study is comparatively large, comprising 27,041 OTUs across 3,849 samples. To test whether the observed trends were robust to dataset scope, we conducted two downsampling experiments (Figure 5). First, we randomly selected between 5-50 samples per body site (25-250 samples in total), re-calculated co-occurrence and phylogenetic networks and assessed body site separation by all 11 indices, at 10 iterations per down-sampling step (Figure 5A; see Figure S3A for the corresponding plot on the R^2^ statistic). We found that even for the smallest tested dataset, TINA and PINA indices provided much clearer partitioning by body site, although ranks by partitioning effect sizes varied across down-sampling iterations. Next, we randomly selected 1,000 samples from which we drew 1,000 sequences each and down-sampled these to 50 sequences per sample in several steps, at 10 iterations per step, recalculated co-occurrence and phylogenetic networks and quantified community similarity (Figures 5B and S3B). Likewise, we observed that TINA and PINA provided much better separation by body site than all other indices, even at a drastically small size of 50 sequences per sample.

**Figure 5.**
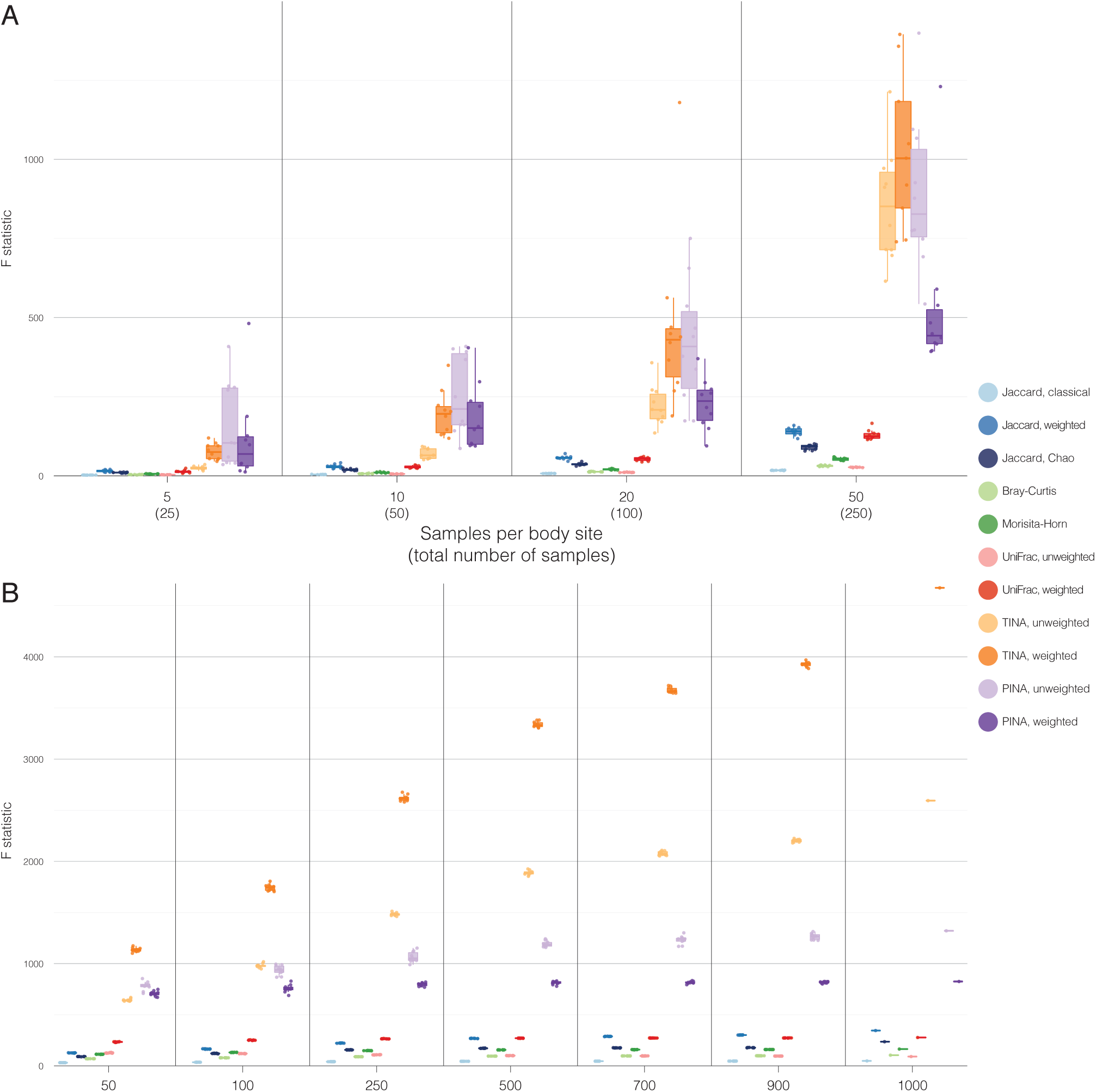
TINA and PINA detect strong partitioning by body habitat even for very small datasets. Adonis F statistic for separation by body site according to all 11 tested indices, for datasets randomly down-sampled following two different regimes. Taxa co-occurrences and phylogenetic interactions were re-calculated for every down-sampled dataset. (A) 5, 10, 20 and 50 samples per body site were randomly selected from the full HMP dataset, at 10 iterations per down-sampling step. (B) 1,000 randomly selected samples were down-sampled to a depth of 1,000 sequences per sample; this dataset was then further down-sampled to 50 sequences per sample in several steps, at 10 random iterations per step. Figure S3 shows corresponding plots on R^2^ values.

### Biogeographical and physicochemical gradients structuring oceanic micro-eukaryote plankton communities are best captured by TINA

The TARA Oceans project has provided a very rich and multifaceted census of the world’s oceans along several geographical and physicochemical gradients. We re-analysed TARA data on micro-eukaryote plankton diversity in the sunlit ocean ^35^ to test the performance and versatility of TINA and other indices at representing different ecological signals. The dataset contained 77 samples from 43 stations along a wide geographical gradient (as shown in Figure 6A), taken at two depth levels, subsurface water (SUR) and deep chlorophyll maximum (DCM), and with body size filters ranging from 0.8-2,000μm. We calculated pairwise community distances according to JCI, JCW, JCC, BC, MH, TU and TW.

**Figure 6.**
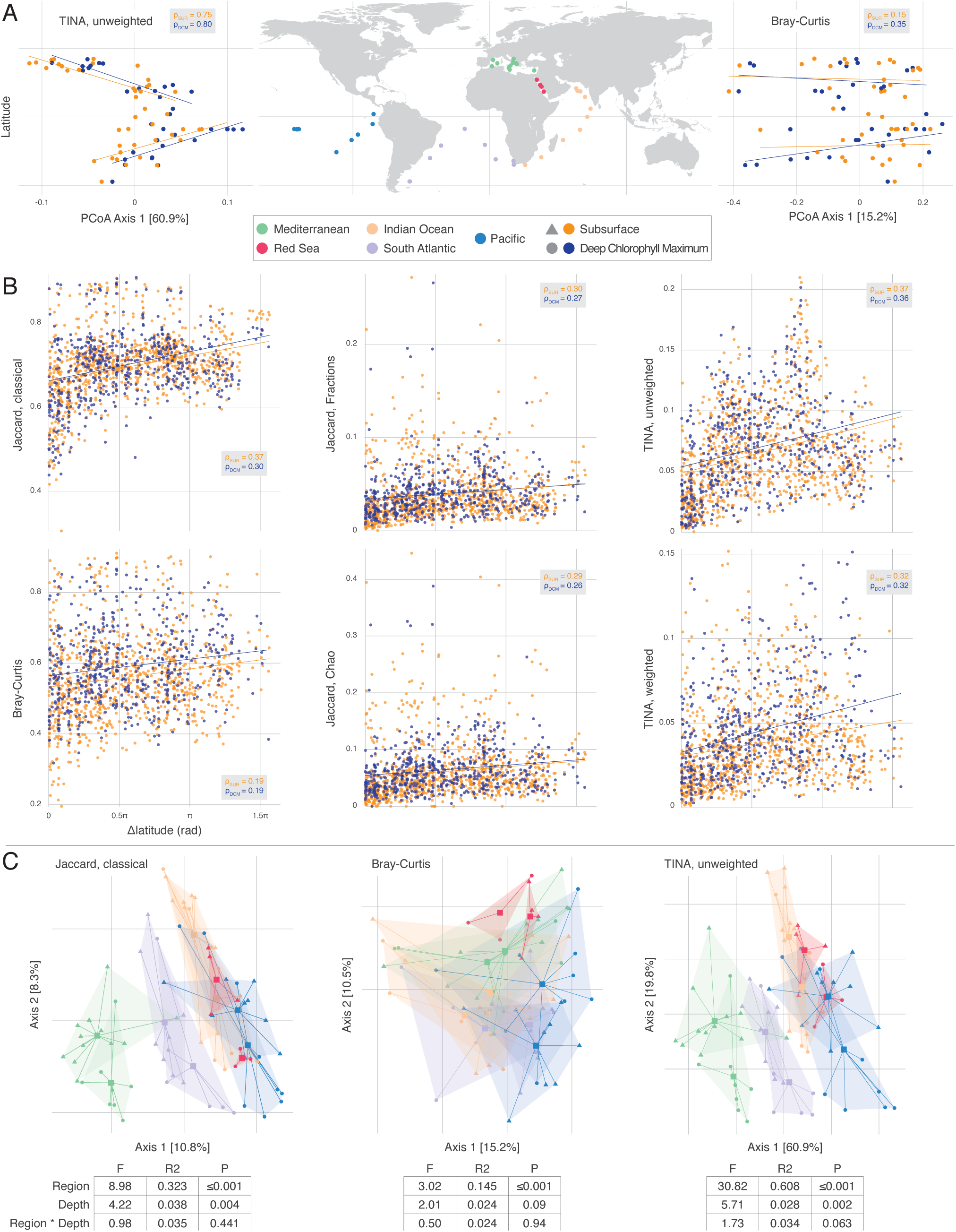
TINA captures biogeographical and physicochemical trends. (A) TARA Oceans sampling locations, coloured by assigned oceanic world region (middle). The first axes of PCoA ordinations on TINA (left) and Bray-Curtis (right) dissimilarities correlated differentially well with latitude both for subsurface (SUR, orange) and deep chlorophyll maximum (DCM, blue) samples, both for the northern and southern hemisphere. (B) Spearman correlations of six different community distances against absolute difference in latitude (Δlat in rad). (C) PCoA ordinations and PERMANOVA statistics on dataset partitioning by factors region, depth and the region*depth interaction term for JCI, BC and TU.

To test whether community similarity follows a latitudinal gradient, we correlated the first component of a Principal Coordinates ordination (PCoA) to latitude, separately for the northern and southern hemispheres (Figure 6A). We observed that the dominating component of an unweighted TINA-based PCoA correlated well with latitude, both for SUR (ρ_Spearman_=0.75) and DCM (ρ=0.8) water layers. In contrast, an ordination based on BC (the index used in the original study) showed much weaker correlations for the dominating component (ρ=0.15 and ρ=35, respectively). We tested for these effects more systematically by correlating pairwise community distances to absolute differences in latitude between samples (Figure 5B). While all indices provided positive correlations, TU and JCI showed the strongest trend for SUR samples (ρ_Spearman_=0.37), while TU and TW showed the strongest signals for DCM (ρ=0.36 and ρ=0.32). Similar trends were observed for correlations of community distance to geographical distance (Table S1). Thus, although JCI performed surprisingly well, TINA outperformed other indices at detecting a biogeographical signal. This can probably be rationalized when considering that taxa co-occurrence is expected to be in part determined by geography, which in turn also correlates with many other ecological parameters like water temperature, irradiation, salinity, etc.

Finally, we tested how well pairwise community distances represented the factors sampling region and depth; Figure 6C shows PCoA ordinations and PERMANOVA statistics for JCI, BC and TU. While all three indices provided significant separation by sampling region, water layer effects were significant only in JCI and TU, while only TU detected any effect of the region*depth interaction term. Moreover, effect sizes as expressed in F statistics and total variance explained by these three factors (summed R^2^) were considerably higher for TU than for JCI and BC, indicating a much more pronounced partitioning according to these terms.

## Discussion

The question of what processes are rendering communities similar or dissimilar to each other is fundamental to the study of diversity. Traditionally, community similarity has been quantified from compositional overlap, considering taxa as independent of each other and describing community structure based on census data. More recently, measures based on additional signals have become increasingly popular, most prominently phylogenetic indices based on shared evolutionary history. Such approaches take into account relationships between taxa, under the assumption that phylogenetic relatedness implies ecological and functional similarity. In this study, we have introduced a set of indices that follow a different rationale: we propose to quantify community similarity in terms of interactions between taxa.

There are several arguments for doing so. First, taxa interactions are a fundamental ecological parameter, at the heart of community ecology: they are important drivers of community assembly, composition and dynamics. Second, it is therefore meaningful to characterise taxa and their relationships based on their interactions: two taxa that are highly similar in terms of their interactions with other taxa can be considered ecologically almost equivalent, they can be assumed to fulfill similar roles in a community. Our approach captures this signal: we argue that it is meaningful to consider communities as similar which contain ecologically similar taxa. Third, interaction network analysis has proven to be a very powerful tool in unravelling complex community dynamics, but its findings are often difficult to connect to the level of community-level diversity patterns. Our approach may help to bridge this analysis gap, by providing diversity indices which are based on networks and can be interpreted in a network framework. For example, TINA essentially quantifies community distance as the distance on a taxa co-occurrence network. Finally, our approach is versatile: there are many types of ecological interactions, at many different levels, and in principle, interaction-adjusted indices like TINA and PINA can be formulated for all of them.

In this study, we have focused on two specific types of interaction-adjusted indices, TINA and PINA, and benchmarked their performance in a re-analysis of two large and complex datasets. TINA by far outperformed all other indices at discriminating human body habitat-specific communities (Figure 2), even for very small datasets (Figure 5). Moreover, TINA best captured biogeographical trends and partitioning by the factor ‘region’ and ‘water depth’ for microeukaryotic plankton communities (Figure 6). These results can be interpreted in light of how TINA is computed. Taxa co-occurrence networks, on which TINA is based, capture an “integrated” ecological association signal, in quantifying the observable outcome of the interplay of different levels of taxa interactions as patterns of co-abundance and avoidance ^18^. Thus, they can reveal network structures that are specific to a given type of habitat (as shown e.g. in Figure 3) or structured according to their response to an ecological gradient or perturbance. Figure 4 illustrates anecdotally how TINA can capture such signals to assign community similarities that are more in line with ecological reality than count-based indices. The fact that TINA provides such strong separations of body habitats and strong correlations with biogeography and other factors for plankton communities means that these factors have a strong and specific influence on taxa co-occurrence; subsequent diversity analyses should take this into account and interpret accordingly.

Likewise, PINA and other phylogenetic indices such as UniFrac are based on phylogenetic relatedness, operating under the assumption that shared phylogeny implies not only a shared evolutionary history, but similar ecological roles and functional profiles. However, by definition, an inference of ecological similarity from phylogeny is necessarily indirect and imperfect, although it is arguably a valid enough assumption for many phylogenetic clades. In all relevant tests, unweighted and/or weighted PINA, based on phylogenetic “interaction” networks, outperformed UniFrac, based on shared phylogeny only, even for very small datasets. Interestingly, unweighted PINA and UniFrac were the only indices to detect partitioning by urogenital habitats (Figure 2), a task at which almost all count-based indices, as well as TINA, failed. As discussed above, this is likely due to differential factors and mechanisms shaping community structure in urogenital habitats, compared to other habitats. However, the fact that we did observe differential trends in index performance emphasises the importance of a multifaceted approach: by applying count-based, phylogenetic and interaction-adjusted indices, different aspects of community similarity are quantified which can be interpreted in context of each other, with the potential to reveal biological insights beyond the scope of mono-dimensional approaches.

One possible drawback of interaction-adjusted indices is that they are not context-invariant: their values will always depend on analysis scope and on the system under consideration, as they vary with varying network structure. While count-based indices will always assume the same similarity for two communities, independently of the remaining dataset, interaction-adjusted indices may change asymptotically when more data is added, simply due to (subtle) changes in network structure. However, this behaviour can indeed also be an asset, for example when comparing multiple datasets in a meta-study. In such a setup, “globally” constructed networks may mitigate dataset-specific noise, introduced e.g. by sampling methods or limited depth. Likewise, interaction-adjusted indices are not limited to capturing static network architectures, but their flexibility allows to account for conditionally variable networks which are rewired e.g. over time, gradients or in response to specific changing factors.

In the large arsenal of measures for community similarity and, more generally, β diversity, our proposed family of interaction-adjusted indices provide an important, powerful and versatile alternative. By taking taxa interactions into account, they quantify novel aspects of “diversity”, at the very core of community ecology, and may guide biological interpretation of diversity patterns in novel ways.

## Acknowledgements

We would like to thank Janko Tackmann, Jordi Bascompte and Bernhard Schmid of the University of Zurich for helpful discussions during the preparation of this manuscript.

## Author contributions

TSBS performed experiments; TSBS, JFMR & CvM conceived the study and analysed the data; TSBS wrote the first draft of the manuscript and all authors contributed to revisions.

## Supporting Information

**Table S1. Spearman correlations of community similarities to geographical distance for TARA samples**.

**Figure S1. Transformation of taxa co-occurrence into co-occurrence similarity smoothens and sharpens interaction information**.

Heatmaps of correlation strengths are shown for 300 randomly selected OTUs from the set shown in the network of Figure 3, for SparCC co-occurrences (referred to as matrix I_C_ in the Methods section) and transformed SparCC correlations (matrix C). OTUs were clustered by SparCC correlation. Note that the color scale is [-1, 1] for a more intuitive visualisation, while we use a linear transformation to [0, 1] in our analyses.

**Figure S2. HMP taxa co-occurrence network, annotated by taxonomy**.

The same network as in Figure 3 of the main text, with nodes are coloured by OTU phylum-level consensus taxonomy.

**Figure S3. Down-sampling effects on body habitat partitioning, quantified as R^2^**

This figure is analogous to Figure 5 of the main text, but showing down-sampling effects in terms of R^2^ instead of F statistics.

